# Characterization and Optimization of *Streptomyces albidoflavus* MD102 as a Heterologous Expression Chassis

**DOI:** 10.64898/2026.02.27.708677

**Authors:** Sean Lee Qiu En, Guanglei Ma, Hartono Candra, Zhao-Xun Liang

## Abstract

We report the isolation and characterization of *Streptomyces albidoflavus* MD102, a strain that can be used as a microbial chassis for the heterologous production of secondary metabolites. This strain, closely related to the widely used *S. albidoflavus* J1074, exhibits a compact genome, exceptional genetic tractability, rapid growth, and susceptibility to antibiotics. Whole-genome sequencing revealed the metabolic capabilities of *S. albidoflavus* MD102, highlighting its versatility in supporting the production of diverse secondary metabolites. Employing CRISPR/Cas9-assisted genome editing tools, we created mutant strains with reduced genome and cleaner chromatographic background. In addition to the deletion of several biosynthetic gene clusters (BGC), we inserted the global regulator *bldA* gene and geranyl diphosphate synthase (*gpps*) genes and an additional ΦBT1-attB attachment site into the chromosome to enhance the strain’s capability in producing secondary metabolites. *S. albidoflavus* MD102 will be a new addition to the repertoire of existing *Streptomyces* chassis, contributing to the advancement of secondary metabolite discovery and synthetic microbiology.

**IMPORTANCE:** The pursuit of a universal *Streptomyces* microbial chassis for the heterologous production of secondary metabolites has proven elusive, prompting a more pragmatic approach to develop a suite of *Streptomyces* chassis. The current study introduces *Streptomyces albidoflavus* MD102 as a promising heterologous chassis with rapid growth, susceptibility to common antibiotics, and genetic tractability. Its close phylogenetic relation with the widely used versatile *S. albidoflavus* J1074 chassis and the traits gained from strain improvement place the engineered *S. albidoflavus* MD102 strains as useful chassis for the heterologous production of microbial secondary metabolites. A notable feature of *S. albidoflavus* MD102 that distinguishes it from J1074 and other *Streptomyces* chassis is the presence of metabolic genes in its genome putatively responsible for the degradation of aromatic compounds. This characteristic may endow the strain with the capability to convert petrogenic polycyclic aromatic hydrocarbons (PAHs) and substituted aromatics into valuable secondary metabolites.

## INTRODUCTION

Actinomycetes are highly prolific microbial producers of secondary metabolites synthesized by the enzymes encoded by biosynthetic gene clusters (BGCs) (Lee et al., 2020). The abundance of BGCs and the widespread distribution of actinomycete strains in the environments and human digestive tracts suggest the underexplored potential of discovering secondary metabolites with novel structures and bioactivity (Lacey and Rutledge, 2022; Nepal and Wang, 2019). Many cryptic BGCs remain dormant under typical laboratory culture conditions due to repressed gene expression and other inhibitory factors (Liu et al., 2021). Activating silent BGCs to disclose their cryptic secondary metabolites is often challenging, with many reported successes owing to serendipitous discoveries resulting from screening elicitors or fermentation conditions (Bode et al., 2002; Craney et al., 2012; Xia et al., 2020).

One approach that has been explored to access the products of silent cryptic BGCs involves the expression of silent cryptic BGCs in a genetically manipulable and fast-growing heterologous chassis (Ahmed et al., 2020). Several actinomycetes strains from the *Streptomyces* genus, including *S. coelicolor, S. lividans, S. avermitilis, S. albus, S. chattanoogensis, S. griseofuscus, S. atratus, S. venezuelae*, have been rendered to genome engineering and chassis optimization (Ahmed et al., 2020; Gomez-Escribano and Bibb, 2011; Hwang et al., 2021; Liu et al., 2016; Myronovskyi et al., 2018; Whitford et al., 2021; Yang et al., 2022). Among the *Streptomyces* chassis, *S. albidoflavus* J1074 has emerged as a robust and versatile chassis employed by many academic researchers. *S. albidoflavus* J1074 (formerly known as *S. albus* J1074) is a derivative of the *S. albus* strain G that lacks the SalI restriction endonuclease gene. *S. albidoflavus* J1074 has been employed to express BGCs from both *Streptomyces* and non-*Streptomyces* strains such as *Micromonospora*, as demonstrated by the production of thiocoraline, tetarimycin A, Clareposxcins, landepoxins, and other secondary metabolites (Kang and Kim, 2021; Myronovskyi and Luzhetskyy, 2019). Myronovskyi et al created the *S. albidoflavus* J1074 Del14 mutant strain by deleting 15 BGCs from the chromosome of *S. albidoflavus* J1074 (Myronovskyi et al., 2018). The Del14 strain was further modified by introducing additional phage phiC31 *attB* sites to improve the production yield of exogenous secondary metabolites (Myronovskyi et al., 2018). Studies demonstrated that the engineered strains not only exhibited higher fermentation titre but also produced secondary metabolites coded by silent BGCs (e.g., pyridnopyrone, salicylic acid, fralnimycin bhimamycin A and aloesaponarin II). (Fazal et al., 2020; Li et al., 2021; Myronovskyi et al., 2018)

Here we report the isolation and characterization and engineering of *Streptomyces albidoflavus* MD102 as a microbial chassis for the expression of BGCs. *S. albidoflavus* MD102 is closely related to the established *S. albidoflavus* J1074 phylogenetically. *S. albidoflavus* MD102 displays rapid growth and efficient sporulation on standard solid and liquid culture media. Its susceptibility to commonly used antibiotics in research laboratories and natural competence allows easy transformation via *E. coli-Streptomyces* conjugation. Using CRISPR-Cas9 tools for genome editing, we successfully eliminated competing biosynthetic gene clusters (BGCs) and introduced genes to improve the strain for heterologous expression of BGCs. With some unique genomic features, *S. albidoflavus* MD102 stands as a valuable supplement to the repertoire of *Streptomyces* chassis, offering the potential for utilization in heterologous expression and contributing to progress in synthetic biology and biotechnological research.

## RESULTS AND DISCUSSION

### Isolation and taxonomic classification of *Streptomyces albidoflavus* MD102

We recently isolated a collection of actinomycete strains from local soil and aquatic environments in Singapore from which novel secondary biosynthetic pathways have been discovered (Candra et al., 2022; Lee et al., 2024; Low et al., 2020, 2018; Ma et al., 2022). Among them, strain *S. albidoflavus* MD102, initially isolated from a sediment sample obtained from the Sungei Buloh Wetland Reserve, exhibited notable rate of growth and sporulation, surpassing several commonly used *Streptomyces* chassis, including *Streptomyces* chassis, *S. coelicolor* M1154 and *S. lividans* TK24 (Figure 1, Table S1). Initial 16S rRNA sequencing analysis suggested its close relationship with *Streptomyces violascens* ISP 5183, with a 100% similarity at 93.8% coverage according to the EzBioCloud database. However, recognizing the limitations of small subunit rRNA gene sequences for *Streptomyces* spp. phylogenetic classifications, we adopted a whole-genome approach to attain species-level resolution of MD102’s phylogenetic relationship. Utilizing a PacBio and Illumina hybrid approach, whole genome sequencing produced a total length of 7,047,131 bp and a GC content of 73.3% (Figure 2A). Phylogenetic relationships were established by constructing a Genome BLAST Distance Phylogeny (GBDP) tree using the Type Strain Genome Software (TYGS). MD102 clustered within a monophylogenetic clade alongside *S. albidoflavus* J1074, *S. coelicolor* DSM 40233, *S. sampsonii* NBRC 13083, *S. limosus* NBRC 12790, *S. albidoflavus* DSM 40455, and *S. albidoflavus* NRRL B-1271 (Figure 2D). Notably, *S. albidoflavus* J1074 exhibited the highest GBDP score with MD102, registering d_0_, d_4_, and d_6_ scores of 92.5, 90.1, and 94.4, respectively, with a 0.0% difference in GC content. Further confirming this relationship, the OrthoANIu value of 98.81% obtained from the EzBioCloud database in the MD102 and *S. albidoflavus* J1074 comparison surpassed the ANI threshold value of 95%, leading to the classification and naming of the strain as *S. albidoflavus* MD102.

**Figure 1.**
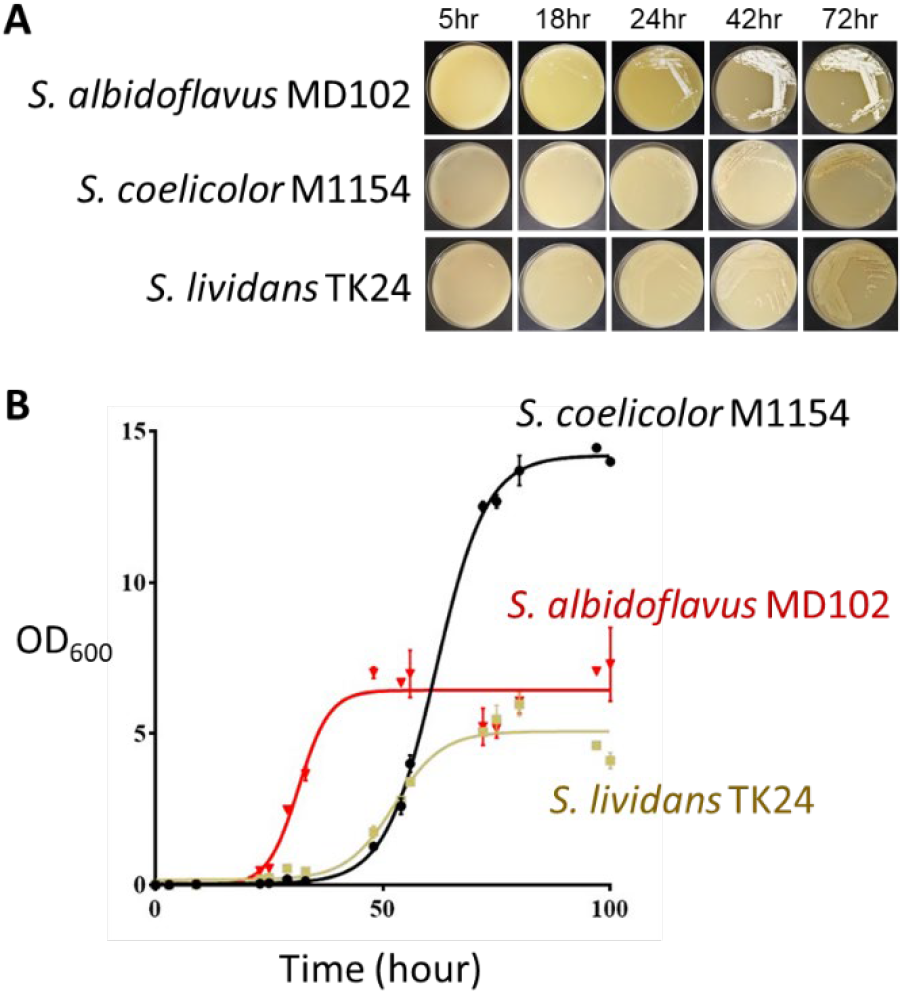
Comparison of the growth of *S. albidoflavus* MD102 to two commonly used *Streptomyces* hosts. A. solid medium. B. CGM liquid medium (30°C).

**Figure 2.**
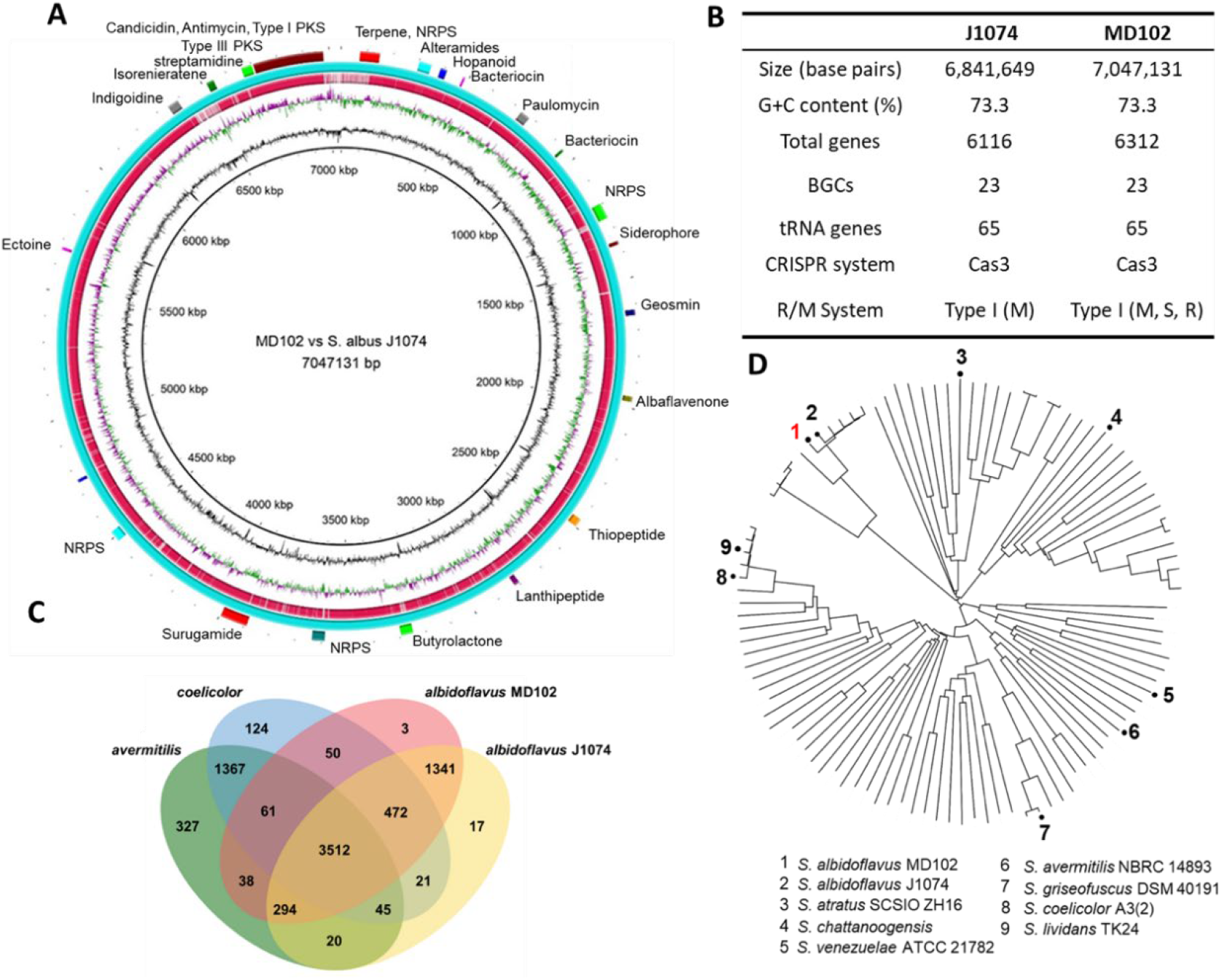
Phylogenetic and genomic characterization of *S. albidoflavus* MD102. (A) Chromosomal genome map of *S. albidoflavus* MD102 with comparisons against *S. albidoflavus* J1074. Tracks from outermost circle to the center: (i) Biosynthetic gene clusters of *S. albidoflavus* MD102 predicted by antiSMASH 7.0.0. (ii) Predicted coding DNA sequence (CDS) of *S. albidoflavus* MD102 in blue. (iii) Predicted coding DNA sequence (CDS) of *S. albidoflavus* J1074 in red. (vi) GC skew of *S. albidoflavus* MD102, the inward facing portion in green indicating higher than average GC skew, and the outward portion in purple indicating below average GC skew. (v) GC content of *S. albidoflavus* MD102, the outward facing portion indicating higher than genome-wide average GC content, whilst the inward portion indicating lower than average GC content. (vi) Marker for for genome size. The genome figure was constructed using DNAPlotter v 18.1.0. (B) Genome features of *S. albidoflavus* J1074 and *S. albidoflavus* MD102.(C) Venn diagram of unique and shared orthologous gene clusters amongst *S. albidoflavus* MD102, *S. albidoflavus* J1074 (CP004370.1), *S. avermitilis* NBRC 14893 (NC_003155), and *S. coelicolor* A3(2) (CP042324). (D) Phylogenetic relationship between *S. albidoflavus* MD102 and commonly used *Streptomyces* chassis.

### Genomic features of *S. albidoflavus* MD102

*S. albidoflavus* J1074 is one of the most used heterologous chassis known for its fast growth and natural competence. Like the fast-growing *S. venezuelae, S. albidoflavus* J1074 contains seven rRNA operons (16S-23S-5S) which may explain the fast growth rate of these two commonly used heterologous hosts (Zaburannyi et al., 2014). Its small genome of 6,841,649 bp also has one of the highest GC content (73.3%) among *Streptomyces* (Zaburannyi et al., 2014). Compared to *S. albidoflavus* J1074, *S. albidoflavus* MD102 has a slightly larger genome (7,047,131 bp) and almost identical GC content (Figure 2B). Genome annotation of *S. albidoflavus* MD102 using RAST identified 65 tRNA genes and seven rRNA operons, the same as *S. albidoflavus* J1074. Like *S. albidoflavus* J1074, *S. albidoflavus* MD102 lacks the SalI restriction modification system that facilitates genetic transformation. However, it possesses the restriction subunit R of the Type I restriction modification system, which may interfere with its ability to accept exogenous DNA.

RAST annotation led to the identification of 6,312 protein coding ORFs, slightly more than the 6,116 found in *S. albidoflavus* J1074. OrthoVenn3 was utilized to compare orthologous proteins of *S. albidoflavus* MD102 with *S. albidoflavus* J1074, *S. coelicolor* and *S. avermitilis*. A total of 5,771 orthologous proteins were identified in *S. albidoflavus* MD102 by OrthoVenn3, with 1,341 uniquely shared with *S. albidoflavus* J1074, whilst only 50 with *S. coelicolor* and 38 with *S. avermitilis*, which agrees with their phylogenetic relationship (Figure 2C). In comparison with the other three model strains, *S. albidoflavus* MD102 has a set of strain-specific genes encoding clusters of uncharacterized proteins. Many of the unique genes are found in the terminal regions of the chromosome, which indicates that those genes are likely non-essential genes acquired later as a result of adaptive evolution.

Among the unique genes found in *S. albidoflavus* MD102 but not in *S. albidoflavus* J1074 are several genes involved in aromatic degradation including biphenyl dioxygenases and other aromatic dioxygenases (Table S2). This might be attributed to the prevalence of polycyclic aromatic hydrocarbons (PAHs) in the mangrove and coastal sediments near the Johor Strait from where *S. albidoflavus* MD102 was isolated. Like other parts of the Singapore coast, the water of Johor Strait ports is heavily polluted due to heavy shipping and petroleum refinery activities (Basheer et al., 2003). We speculated that the enzymes encoded by those genes may enable the strain to grow on aromatic substrates such as biphenyl, toluene, and phenols. Under our lab culturing conditions, *S. albidoflavus* MD102 did not grow on biphenyl as the sole carbon source in our laboratory (data not shown). This may be attributed to the repressed expression of the metabolic genes under our culturing conditions or biphenyl being the wrong substrate. Considering that microbes harbouring similar metabolic genes are known to degrade aromatic compounds (Mahjoubi et al., 2019; Segura et al., 2021), further engineering to activate the expression of the metabolic genes could transform *S. albidoflavus* MD102 into a unique heterologous chassis for the conversion of petrogenic polycyclic aromatic hydrocarbons (PAHs), and substituted aromatics into valuable secondary metabolites.

### The biosynthetic capability of *S. albidoflavus* MD102

We mined the biosynthetic gene clusters (BGCs) of *S. albidoflavus* MD102 using antiSMASH to assess the biosynthetic capacity of *S. albidoflavus* MD102. The AntiSMASH analysis identified 23 Biosynthetic gene clusters (BGCs), the same number as *S. albidoflavus* J1074. However, a close inspection of the BGCs revealed that two of the BGCs are different between the two strains. In *S. albidoflavus* MD102, BGC1 includes genes that encode terpene synthase and NRPS-like biosynthetic enzymes and BGC13 contains polyketide synthase (PKS) genes (Table S4). In *S. albidoflavus* J1074, BGC1 codes for a T1PKs-NRPS hybrid whereas BGC12 codes for a ribosomally synthesized and post-translationally modified peptides (RiPP) pathway.

We cultured *S. albidoflavus* MD102 in various culture media to identify constitutively active BGCs. High-performance liquid chromatography (HPLC) and liquid chromatography-coupled mass spectrometry (LC/MS)-based metabolite profiling suggested the production of several metabolites that include the polyketide-derived secondary metabolites alteramides, paulomycin, surugamide, candicidin, and the polyketide-peptide hybrid antimycin. The production of the antifungal polyene compound candicidin is consistent with the observation that *S. albidoflavus* MD102 displayed strong antifungal activity against *Candida albicans*, with a large zone of inhibition on antifungal assay plate overlay (date not shown). The results from the AntiSMASH mining analysis underscore the potential of *S. albidoflavus* MD102 to synthesize a diversity of secondary metabolites, indicating that its genetic makeup encompasses a robust metabolic network conducive to the heterologous production of secondary metabolites. Considering that only a small fraction of the predicted secondary metabolites has been identified by the metabolite profiling, many of the endogenous BGCs are either silent or expressed at low levels under laboratory fermentation conditions. The discrepancy between the large number of secondary metabolites suggested by AntiSMASH analysis and the small number of metabolites detected underpins the highly regulated production of secondary metabolites in *S. albidoflavus* MD102 as observed for other *Streptomyces*.

### Strain improvement by genome editing

We employed the CRISPR/Cas9 tool for rational genome editing in an effort to further improve *S. albidoflavus* MD102 as a heterologous host (Table S3). In comparison to the RedET double crossover methodology utilized for BGC deletion in *S. albidoflavus* J1074, the CRISPR/Cas9 method proves more efficient and avoids leaving scars in the chromosome. Our focus was on the targeted deletion of constitutively active BGCs, including alteramide, paulomycin, BGC16 (NRPS), candicidin, and antimycins. These deletions aimed to simplify the background of the HPLC chromatogram during metabolite profiling while boosting the supply of precursors (e.g., amino acids, acetyl-CoA, malonyl-CoA) and cofactors (e.g., NADPH). The clean chromatogram background would facilitate the detection of compounds and simplify the compound-isolation process. Meanwhile, we also constructed the CRISPR/Cas9 plasmids in such a way that we could concurrently insert exogenous genes into the chromosome to enhance heterologous expression and BGC activation (Figure 3A). The inserted genes include the global regulator *bldA* gene sourced from *S. coelicolor*, encoding tRNAleu (UAA). This rare codon UAA is known to constrain the expression of biosynthetic enzymes in *Streptomyces*. The integration of *bldA*, coupled with the constitutively.active promoter SF14p, is anticipated to facilitate the translation of biosynthetic enzymes. Another gene we planned to knock in is *gpps* (from *S. tasikensis* P46 (Candra et al., 2022; Ma et al., 2022) that encodes a geranyl diphosphate synthase that synthesizes geranyl pyrophosphate (GPP), which is the precursor for the biosynthesis of diverse terpenes and terpenoids. The introduction of *gpps* under the control of the constitutively active promoter KasOp aims to augment the cellular supply of GPP, enhancing the production of terpene-based natural products. We also integrated an additional attachment site, ΦBT1-attB, into the BGC23a region. This novel attachment site differs from the ΦC31-attB site present in *S. albidoflavus* MD102, enabling the integration of biosynthetic genes into two distinct sites.

**Figure 3.**
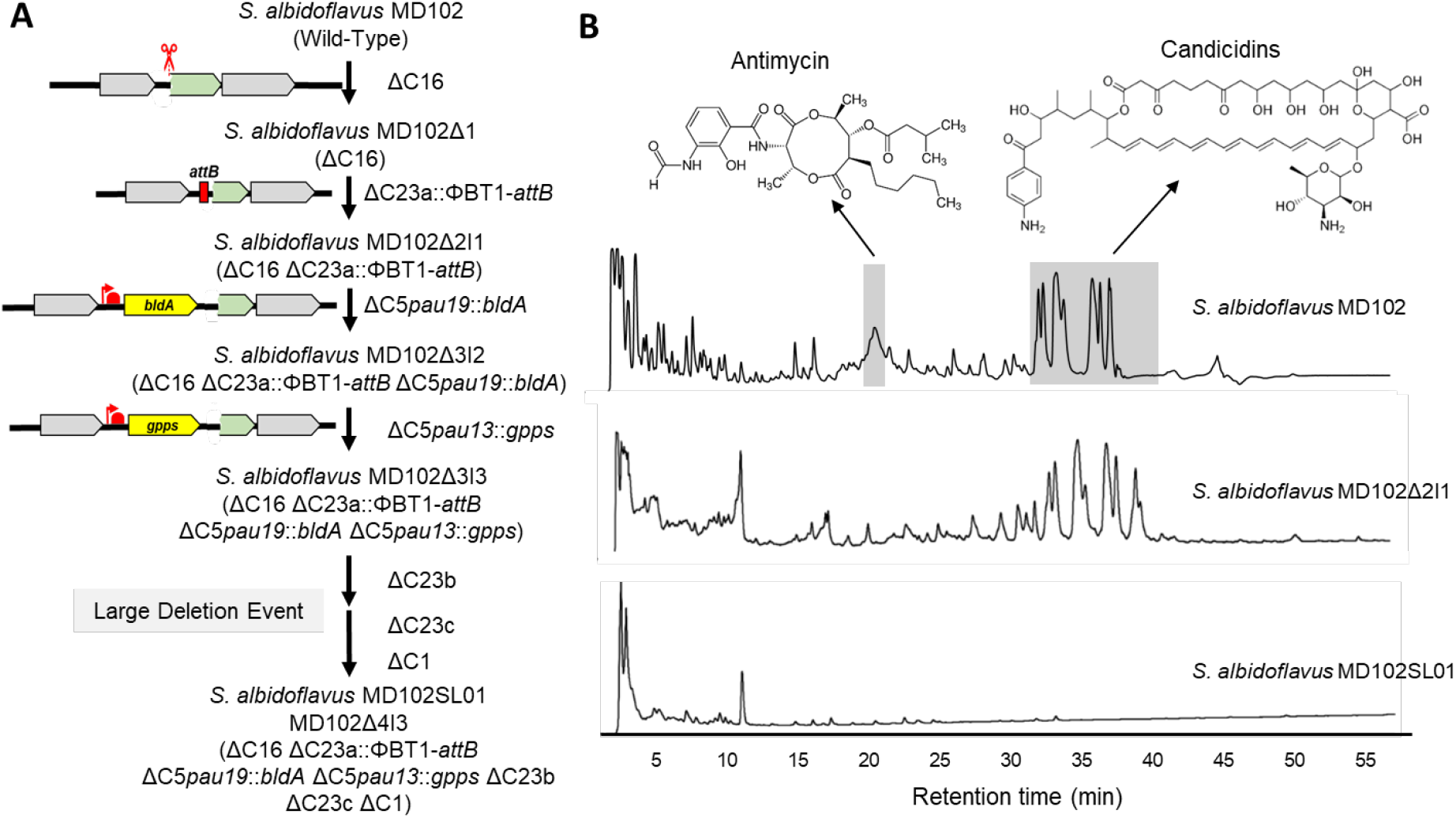
Production of secondary metabolites by *S. albidoflavus* MD102 wild-type and mutant strains. (A) Pedigree of *S. albidoflavus* MD102 mutant strains. Following several rounds of CRISPR-Cas9 facilitated genome editing and an unexpected large deletion event, four endogenous secondary metabolite gene clusters were deleted, and *bldA,gpps*, and ΦBT1-attB were introduced into the genome. BGC16 (as denotated as C16 was initially deleted), followed by candicidin BGC (C23a) deletion and ΦBT1-attB insertion into the deleted region, paulomycin BGC (C5) deletion, and *bldA* and *gpps* insertion into the deleted region, and finally a large deletion event resulting in deletion of antimycin BGC (C23b), BGC23c (C23c), and BGC1 (C1). (B) HPLC analysis confirmed the loss of antimycin and candicidins production in *S. albidoflavus* MD102 mutant strains.

For the genome-editing experiments, the temperature-sensitive pSG5-based pCRISPR-Cas9 plasmids containing the relevant homologous arms and sgRNA sequences were introduced into *S. albidoflavus* MD102 following protocols we optimized previously in our laboratory (Candra et al., 2022; Lee et al., 2024; Low et al., 2020, 2018; Ma et al., 2022). The culmination of the genome editing process, marked by the deletion of the antimycin and other constitutively expressed BGCs, led to a substantially simplified HPLC profile for the resulting mutant strain *S. albidoflavus* MD102SL01 (Figure 3B). Illumina-based whole genome sequencing was conducted to validate the integrity of the MD102SL01 genome after several rounds of sequential gene deletion and insertion. Genome sequencing affirmed the successful deletion of paulomycin (BGC5), BGC16-NRPS, Candicidin (BGC23a), and antimycin (BGC23b) BGCs, as well as the insertion of *bldA, gpps*, and ΦBT1-attB, thereby confirming the efficacy of the CRISPR-Cas9 tool in genome editing. However, an unforeseen deletion of the 5’ and 3’ chromosome end regions was detected. At the 5’ end, a loss of 154,300 bp occurred, and at the 3’ end, a loss of 292,451 bp transpired, resulting in a total loss of 446,751 bp. This unexpected deletion resulted in the additional loss of BGC1 and BGC23 clusters. This substantial deletion event was likely an outcome of the BGC23b (antimycin BGC) deletion during the fifth round of CRISPR-Cas9 editing. Together, the results demonstrate the compatibility of *S. albidoflavus* MD102 with CRISPR-Cas9-based genome editing techniques. The occurrence of this unexpected chromosomal deletion is not an uncommon phenomenon, as documented in our previous studies and those conducted by others (Hoff et al., 2018; Lee et al., 2024).These findings underscore the importance of carefully choosing the PAM and Cas9-cutting sites to mitigate the risk of inadvertent homologous recombination events in *S. albidoflavus* MD102 and other Streptomyces strains.

### Assessment of genetic tools for heterologous expression in *S. albidoflavus* MD102

We further assessed the natural competence of *S. albidoflavus* MD102 using commonly employed vectors by academic researchers, including several conjugative/integrative vectors derived from pIJ101 and pSET152, such as pIJ12551 and pIJ8630. Through *E. coli-Streptomyces* conjugation, these vectors were efficiently introduced into *S. albidoflavus* MD102 and integrated into the chromosome. The vectors derived from pIJ101 and pSET152 rely on the ΦC31 integrase gene for chromosomal integration. Their successful integration suggests that the attB site in the pirin-like gene of *S. albidoflavus* MD102 is active. For the heterologous expression of large BGCs, transformation by plasmids that can carry large DNA fragments of 30-150Kb is crucial. We successfully introduced cosmid-based pIJ10702 vector and BAC-based pSMART-BAC-S into *S. albidoflavus* MD102 via *E. coli-Streptomyces* conjugation. However, attempt with the PAC-based pESAC13A vector was unsuccessful, as the *E. coli-Streptomyces* conjugation did not yield any exconjugants.

Having established the transformation protocols for *S. albidoflavus* MD102, we proceeded to evaluate the strength of several widely used constitutively active promoters. These promoters were cloned upstream of the EGFP gene in the reporter vector pIJ8630 and introduced into *S. albidoflavus* MD102. By quantifying EGFP fluorescence intensity in the culture broth and normalizing against the ermEp* promoter, we observed that KasOP*, gapdhp(EL), and gapdh(KR) promoters induced the highest EGFP expression. In contrast, ermEp* and SF14p induced moderate EGFP expression (Figure 4A). A similar expression pattern was noted in *S. albidoflavus* J1074, reaffirming the comparable cellular environment and genetic background of the two strains.

**Figure 4.**
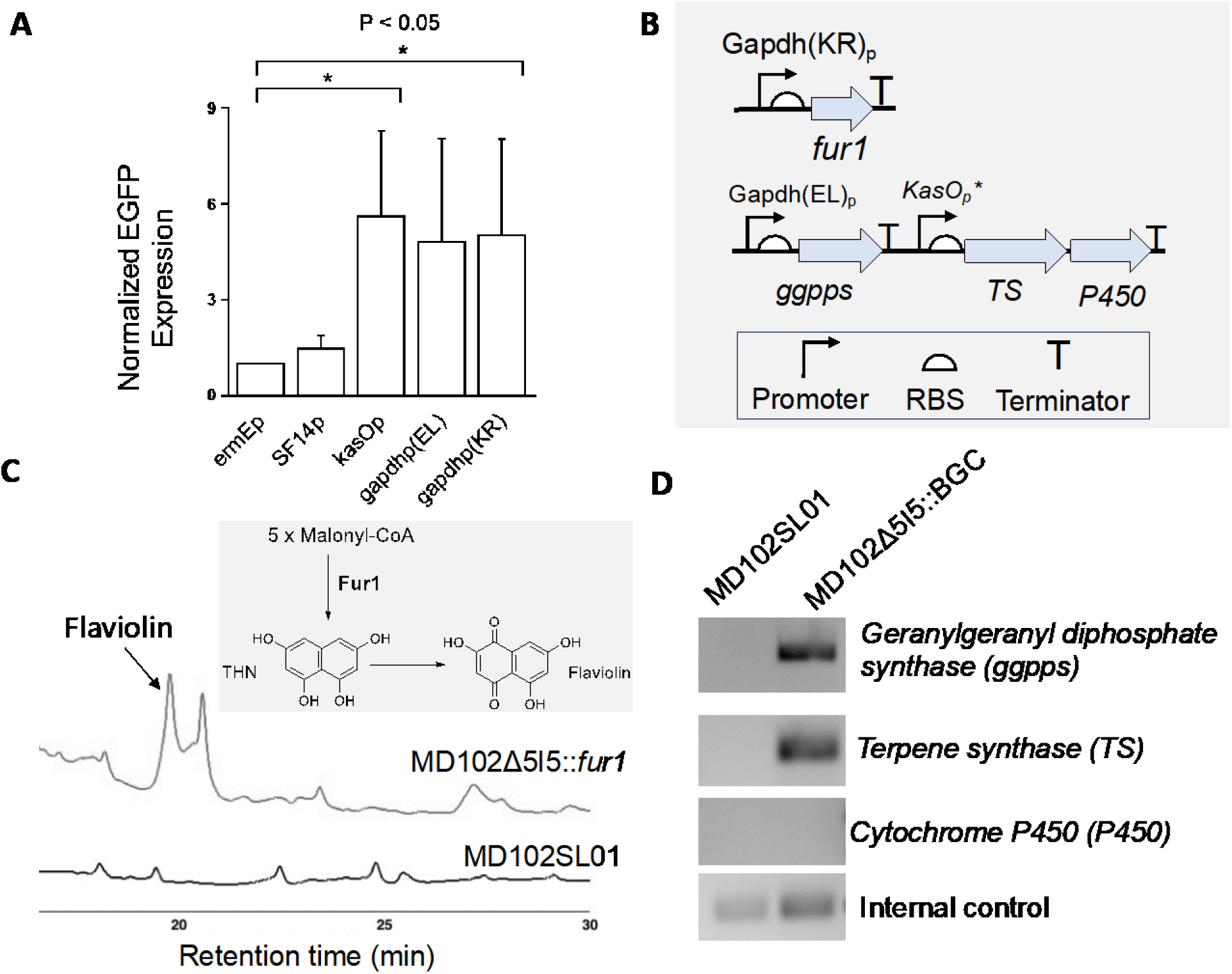
Heterologous production of secondary metabolites using *S. albidoflavus* MD102. (A) Promoter strength comparison in *S. albidoflavus* MD102 using constitutively active promoters and pIJ8630 EGFP reporter vector. EGFP Fluorescence intensities were normalised against ermEp, and the values are means with standard deviation (SD) from three independent experiments. (B) Schematic showing the two exogenous biosynthetic gene circuits integrated into *S. albidoflavus* MD102. (C) HPLC chromatogram showing the production of flaviolin in *S. albidoflavus* MD102 mutant. (D) Transcriptional analysis of the cryptic terpene BGC genes using RT-PCR. *hrdB* housekeeping gene is used as internal control.

Due to the cost and time-intensive nature of cloning large BGCs, an extensive evaluation of *S. albidoflavus* MD102’s ability to express diverse biosynthetic pathways from other actinomycetes was not undertaken in this study. However, given the high similarity between *S. albidoflavus* MD102 and the well-characterized *S. albidoflavus* J1074, we anticipate that *S. albidoflavus* MD102 will exhibit similar behaviour as J1074. Instead of focusing on diverse BGCs, we focused on determining whether *S. albidoflavus* MD102 can effectively express biosynthetic genes under the control of orthogonal or non-native promoters—a critical aspect for the expression of refactored BGCs and engineered pathways.

We first cloned the *fur1* gene from the furaquinocin BGC of *S. tasikensis* P46 and integrated it into the MD102 chromosome using the conjugative/integrative vector pIJ8630, coupled with the robust promoter gapdhp(KR) (Figure 4B).

The *fur1* gene encodes a type III polyketide synthase (PKS) responsible for catalyzing the production of 1,3,6,8-tetrahydroxynaphthalene (THN) from malonyl-CoA molecules, leading to the formation of the red-colored pigment flaviolin. Culturing *S. albidoflavus* MD102SL01::*fur1* in natural product (Table S5). The refactored BGC was created with the three genes of the cryptic BGC the native BGC, the terpene synthase (*ts*) and the cytochrome P450 (*P450*) genes were kept in the same operon without adding an extra promoter for the *P450* gene. While successful integration of the refactored BGC into the MD102SL01 chromosome was achieved via *E. coli*-*Streptomyces* conjugation, the expected diterpenoid product was not detected in the fermentation culture. RT-PCR analysis TSB liquid media resulted in the successful production of the anticipated red-colored pigment, flaviolin, validated using a standard (Figure 4C).

We refactored a cryptic BGC (BGC6) from *S. tasikensis* P46 predicted to produce a diterpenoid equipped with two orthogonal promoters and terminators (Figure 4B). Following the design of revealed expression of the geranylgeranyl pyrophosphate synthetase (*ggpps*) and the *ts* genes, but the mRNA of the cytochrome P450 was not detected. This suggests that the anticipated diterpenoid was not observed likely due to the lack of cytochrome P450 expression or issues with protein translation or solubility. The observations indicate that the genetic vectors and orthogonal promoters are compatible with *S. albidoflavus* MD102. Nonetheless, predicting the expression levels of genes proves challenging, underscoring the necessity for experimental validation to confirm successful gene expression in this microbial chassis. In summary, we propose that *S. albidoflavus* MD102 can be used as a microbial chassis for the heterologous production of secondary metabolites. Sharing key characteristics with its close relative, *S. albidoflavus* J1074, this strain exhibits several traits that render it well-suited for expressing heterologous biosynthetic pathways. *S. albidoflavus* MD102 possesses primary metabolic networks that are expected to support the production of polyketides, non-ribosomal peptides, and other natural products. The accessibility of the comprehensive genome sequence of *S. albidoflavus* MD102 enables strain enhancement through rational genome engineering. The strain displays outstanding genetic tractability, allowing for easy genetic manipulation to enhance its traits through rational approaches. The targeted removal of competing and interfering biosynthetic pathways and a ~200kb region has resulted in mutant strains with reduced genome and cleaner chromatographic background. The incorporation of an additional attB site, along with *bldA* and *gpps* genes, has yielded strains that will facilitate the expression and activation of silent BGCs, as well as enhancing fermentation titer. These engineered mutant strains represent promising starting points for the development of customized chassis tailored for the production of polyketides, terpenes, and various secondary metabolites.

## MATERIALS AND METHODS

### Isolation of actinobacterial strains

1m depth soil sediments were previously obtained from Sungei Buloh Wetland Reserve, Singapore (1°26’41.172”N, 103° 43’36.12”E) via a sterile stainless-steel soil sampler. The samples were desiccated using microwave treatment, and sterile water used in various dilutions to obtain single colonies on various solid agar media.

### Actinobacterial culture conditions

Routine culturing of *Streptomyces* strains was done on mannitol soy (MS) solid media (20 g/L BactoTM agar, 20 g/L mannitol, and 20 g/L soya flour), tryptic soy broth (TSB) (30 g/L TSB powder) (Merck Millipore, Germany) liquid media and GYMose (20 g/L glucose, 10 g/L maltose, 5 g/L yeast extract and 2 g/L CaCO3) liquid media. Selective antibiotics were used at 50 μg/mL apramycin, 25 μg/mL chloramphenicol, 50 μg/mL kanamycin, and 25 μg/mL nalidixic acid. Culturing of *Streptomyces* strains was done at 30°C, and for liquid culture shaking was carried out at 180 rpm.

### Actinobacterial Genomic DNA extraction

The extraction of *Streptomyces* genomic DNA was done using the Hexadecyl trimethyl-ammonium bromide (CTAB) buffer (2 g CTAB, 10 mL 1 M Tris (pH 8), 4 mL 0.5 M ethylenediamine tetra-acetic acid disodium salt (pH 8), 28 mL 5 M NaCl, 1 g polyvinyl pyrrolidone (MW 40 kDa) and 58 mL water) method.

### Phylogenetic and Comparative Genomics analysis

Next-generation whole genome sequencing via Illumina 2x150bp paired-end configuration was used to obtain the genome sequence of *S. albidoflavus* MD102. The establishment of the phylogenetic relationships and closest related strains for *S. albidoflavus* MD102 was carried out using the TYGS server (https://tygs.dsmz.de) and and visualised by the online software iTOL (https://itol.embl.de/). The d_0_, d_4_, and d_6_ scores are defined by formula d_0_: the length of all high-scoring segment pairs (HSP) divided by total genome length, formula d_4_: the sum of all identities found in HSPs divided by overall HSP length, and formula d_6_: the sum of all identities found in HSPs divided by total genome length. The OrthoANIu value was obtained from the ANI calculator from EZBioCloud. RAST (Rapid Annotation using Subsystem Technology) was used to predict the open reading frames and carry out genome annotation. OrthoVenn3 was used to compare orthologous protein clusters. The antiSMASH database was used to identify biosynthetic gene clusters. Unique aromatic degradation genes were blasted against common *Streptomyces* genomes in the NCBI including *Streptomyces coelicolor* A3(2) (taxid:100226), *Streptomyces avermitilis* MA-4680 (taxid:227882), and *Streptomyces albidoflavus* (taxid:1886). These genomes were used as queries for ortholog identification, and results were filtered by 75% sequence identity, 50% coverage, and 1e-10 expected value.

### Fermentation and metabolite profiling

*Streptomyces* strains were first cultured on MS agar. Spores from the MS agar plates were inoculated into 10ml liquid TSB media and cultured at 30°C 180rpm for 4-5 days. Subsequently, this starter culture was used to subculture 50ml of liquid media at 2% and grown for 1 week. Ethyl acetate was used for metabolite extraction of the culture broth while methanol was used for the biomass. HPLC analysis was performed on Agilent1200 HPLC-DAD system equipped with an ODS column (Pursuit XRs: 250 mm × 4.6 mm, 5 µm) using a linear gradient of CH3CN in H2O with 0.1% formic acid (0–5 min, 10%–20% CH3CN; 5–35 min, 20%–70% CH3CN; 35–50 min, 70%–90% CH3CN; 50–60 min, 90%– 100% CH3CN; 60–70 min, 100% CH3CN) at a flow rate of 1.0 mL/min.

### Molecular biology techniques and molecular cloning

PCR reactions were carried out using Q5 High Fidelity Polymerase (NEB) in accordance with the manufacturer’s instructions. The purification of plasmids (Qiagen Spin Miniprep) and gel extractions (Zymo Research Zymoclean Gel DNA Recovery Kit) were done according to the manufacturer’s instructions. The purified DNA was eluted in Milli-Q water (Merck Millipore) or provided elution buffer. DNA digestion was carried out using various NEB restriction enzymes at their recommended optimum temperatures overnight.

### CRISPR/Cas9-assisted genome modification

pCRISPR-Cas9 derivatives containing sgRNA and homology arms were constructed as described by (Tong et al., 2015). In brief, short oligonucleotides corresponding to the sgRNA were cloned into the pCRISPR-Cas9 vector after NcoI and SnaBI digestion. Following which, homologous recombination templates were introduced into the vectors via Gibson assembly after StuI digestion. For conjugation of plasmids including the pCRISPR-Cas9 vector into *Streptomyces, E. coli* ET12567/pUZ8002 strains containing the vector of interest were grown at 37c with 50 μg/μL apramycin, 25 μg/μL chloramphenicol and 50 μg/μL kanamycin. Successful exconjugants were screened on MS plate containing 50 μg/mL apramycin and 25 μg/mL nalidixic acid. Subsequent 5 μg/mL thiostrepton was used for the activation of the tipA inducible promoter on pCRISPR-Cas9. Curing of pCRISPR-Cas9 plasmid was done in TSB liquid culture at 37c for one week.

### Measurement of promoter strength

*Streptomyces* spores were first inoculated into 50ml TSB liquid media starter cultures for two days, before subculturing into 50ml TSB liquid media for another three days with the addition of 30 5-mm glass beads. 10ml of culture was extracted and washed and resuspended with 1x PBS. 100ul of cells from 10ml of 1x PBS cell suspension was used for measurement of OD600 and normalization of cell concentration to 0.5 OD600. Fluorescence measurements were carried out using a Tecan Infinite® 200 PRO Plate Reader. For EGFP, the excitation and emission wavelengths used were 485 nm and 515 nm respectively and readings were done in triplicates for each sample.

### RNA isolation and sequencing

*S. albidoflavus* MD102 was grown in TSB media for 72hrs in baffled flasks. Cells were collected and homogenized using a pestle. 1mg/ml working concentration of lysozyme was used for incubation at 37 ºC for half an hour. Cells were freeze-thawed using liquid nitrogen for 5-10 cycles. TRIzol Reagent was added according to the manufacturer’s instructions (Invitrogen). Finally, total RNA was extracted and purified using the Purelink RNA Mini kit according to manufacturer’s instructions (Thermo Fisher Scientific). DNAse-I (Thermo Fisher Scientific) was added in accordance with the Purelink RNA Mini kit protocol.

### Flaviolin production in *S. albidoflavus* MD102

pIJ8630-*fur1* was conjugated into MD102SL01 as described above. The pink colouration of the colonies was observed when cultured on mannitol soy (MS) solid media. MD102SL01-*fur1* exconjugants were cultured in liquid media (GYMose and TSB) in baffled flasks for 5 days before extraction of metabolites using ethyl acetate and methanol. The production of the metabolite was monitored using HPLC and LCMS.

## Acknowledgement

We are grateful for the generous financial support from NRF, Singapore (NRF-CRP31-0005 grant to Z.X.L) and MOE, Singapore (RG37/23 to Z.X.L). We also thank the contribution of Mr. Melvin Wei Sheng Lim, Dr. Howard Saw, Rachel Andrea Chea Yuen Fong, Jiajun Lee, Danial Wong for their contribution to strain characterization, gene cloning, and genome editing during the early stage of the project.

## Supplementary material

Material and methods, supplementary tables and figures that contain information about the cell lines, DNA material, and analytical data obtained through NMR and mass spectrometry analyses are included in the Supporting Information.

## References

Ahmed, Y., Rebets, Y., Estévez, M.R., Zapp, J., Myronovskyi, M., Luzhetskyy, A., 2020. Engineering of Streptomyces lividans for heterologous expression of secondary metabolite gene clusters. Microb Cell Fact 19, 5. 10.1186/s12934-020-1277-8

Basheer, C., Obbard, J.P., Lee, H.K., 2003. Persistent Organic Pollutants in Singapore’s Coastal Marine Environment: Part II, Sediments. Water, Air, & Soil Pollution 149, 315–325. 10.1023/A:1025673517831

Bode, H.B., Bethe, B., Höfs, R., Zeeck, A., 2002. Big effects from small changes: possible ways to explore nature’s chemical diversity. Chembiochem 3, 619–627. 10.1002/1439-7633(20020703)3:7<619::AID-CBIC619>3.0.CO;2-9

Candra, H., Ma, G.-L., En, S.L.Q., Liang, Z.-X., 2022. Enaminone Formation Drives the Coupling of Biosynthetic Pathways to Generate Cyclic Lipopeptides. ChemBioChem 23, e202200457. 10.1002/cbic.202200457

Craney, A., Ozimok, C., Pimentel-Elardo, S.M., Capretta, A., Nodwell, J.R., 2012. Chemical perturbation of secondary metabolism demonstrates important links to primary metabolism. Chem Biol 19, 1020–1027. 10.1016/j.chembiol.2012.06.013

Fazal, A., Thankachan, D., Harris, E., Seipke, R.F., 2020. A chromatogram-simplified Streptomyces albus host for heterologous production of natural products. Antonie van Leeuwenhoek 113, 511–520. 10.1007/s10482-019-01360-x

Gomez-Escribano, J.P., Bibb, M.J., 2011. Engineering Streptomyces coelicolor for heterologous expression of secondary metabolite gene clusters: Streptomyces host for heterologous expression of gene clusters. Microbial Biotechnology 4, 207–215. 10.1111/j.1751-7915.2010.00219.x

Hwang, S., Lee, Y., Kim, J.H., Kim, G., Kim, H., Kim, W., Cho, S., Palsson, B.O., Cho, B.-K., 2021. Streptomyces as Microbial Chassis for Heterologous Protein Expression. Front. Bioeng. Biotechnol. 9, 804295. 10.3389/fbioe.2021.804295

Kang, H.-S., Kim, E.-S., 2021. Recent advances in heterologous expression of natural product biosynthetic gene clusters in Streptomyces hosts. Current Opinion in Biotechnology, Chemical Biotechnology • Pharmaceutical Biotechnology 69, 118–127. 10.1016/j.copbio.2020.12.016

Lacey, H.J., Rutledge, P.J., 2022. Recently Discovered Secondary Metabolites from Streptomyces Species. Molecules 27, 887. 10.3390/molecules27030887

Lee, N., Hwang, S., Kim, J., Cho, S., Palsson, B., Cho, B.-K., 2020. Mini review: Genome mining approaches for the identification of secondary metabolite biosynthetic gene clusters in Streptomyces. Computational and Structural Biotechnology Journal 18, 1548–1556. 10.1016/j.csbj.2020.06.024

Lee, S.Q.E., Ma, G.-L., Candra, H., Khandelwal, S., Pang, L.M., Low, Z.J., Cheang, Q.W., Liang, Z.-X., 2024. Streptomyces sungeiensis SD3 as a microbial chassis for the heterologous production of secondary metabolites. ACS Synthetic Biology.

Li, X., Hu, X., Sheng, Y., Wang, H., Tao, M., Ou, Y., Deng, Z., Bai, L., Kang, Q., 2021. Adaptive Optimization Boosted the Production of Moenomycin A in the Microbial Chassis Streptomyces albus J1074. ACS Synth. Biol. 10, 2210–2221. 10.1021/acssynbio.1c00094

Liu, Q., Xiao, L., Zhou, Y., Deng, K., Tan, G., Han, Y., Liu, X., Deng, Z., Liu, T., 2016. Development of Streptomyces sp. FR-008 as an emerging chassis. Synthetic and Systems Biotechnology 1, 207–214. 10.1016/j.synbio.2016.07.002

Liu, Z., Zhao, Y., Huang, C., Luo, Y., 2021. Recent Advances in Silent Gene Cluster Activation in Streptomyces. Frontiers in Bioengineering and Biotechnology 9.

Low, Z.J., Pang, L.M., Ding, Y., Cheang, Q.W., Le Mai Hoang, K., Thi Tran, H., Li, J., Liu, X.-W., Kanagasundaram, Y., Yang, L., Liang, Z.-X., 2018. Identification of a biosynthetic gene cluster for the polyene macrolactam sceliphrolactam in a Streptomyces strain isolated from mangrove sediment. Sci Rep 8, 1594. 10.1038/s41598-018-20018-8

Low, Z.J., Xiong, J., Xie, Y., Ma, G.-L., Saw, H., Tran, H.T., Wong, S.L., Pang, L.M., Fong, J., Lu, P., Hu, J.-F., Liang, Y., Miao, Y., Liang, Z.-X., 2020. Discovery, biosynthesis and antifungal mechanism of the polyene-polyol meijiemycin. Chem. Commun. 56, 822–825. 10.1039/C9CC08908J

Ma, G.-L., Candra, H., Pang, L.M., Xiong, J., Ding, Y., Tran, H.T., Low, Z.J., Ye, H., Liu, M., Zheng, J., Fang, M., Cao, B., Liang, Z.-X., 2022. Biosynthesis of Tasikamides via Pathway Coupling and Diazonium-Mediated Hydrazone Formation. J. Am. Chem. Soc. 144, 1622–1633. 10.1021/jacs.1c10369

Mahjoubi, M., Aliyu, H., Cappello, S., Naifer, M., Souissi, Y., Cowan, D.A., Cherif, A., 2019. The genome of Alcaligenes aquatilis strain BU33N: Insights into hydrocarbon degradation capacity. PLOS ONE 14, e0221574. 10.1371/journal.pone.0221574

Myronovskyi, M., Luzhetskyy, A., 2019. Heterologous production of small molecules in the optimized Streptomyces hosts. Natural Product Reports 36, 1281–1294. 10.1039/C9NP00023B

Myronovskyi, M., Rosenkränzer, B., Nadmid, S., Pujic, P., Normand, P., Luzhetskyy, A., 2018. Generation of a cluster-free Streptomyces albus chassis strains for improved heterologous expression of secondary metabolite clusters. Metabolic Engineering 49, 316–324. 10.1016/j.ymben.2018.09.004

Nepal, K.K., Wang, G., 2019. Streptomycetes: Surrogate hosts for the genetic manipulation of biosynthetic gene clusters and production of natural products. Biotechnology Advances 37, 1–20. 10.1016/j.biotechadv.2018.10.003

Segura, A., Udaondo, Z., Molina, L., 2021. PahT regulates carbon fluxes in Novosphingobium sp. HR1a and influences its survival in soil and rhizospheres. Environmental Microbiology 23, 2969–2991. 10.1111/1462-2920.15509

Tong, Y., Charusanti, P., Zhang, L., Weber, T., Lee, S.Y., 2015. CRISPR-Cas9 Based Engineering of Actinomycetal Genomes. ACS Synth Biol 4, 1020–1029. 10.1021/acssynbio.5b00038

Whitford, C.M., Cruz-Morales, P., Keasling, J.D., Weber, T., 2021. The Design-Build-Test-Learn cycle for metabolic engineering of Streptomycetes. Essays in Biochemistry 65, 261–275. 10.1042/EBC20200132

Xia, H., Li, X., Li, Z., Zhan, X., Mao, X., Li, Y., 2020. The Application of Regulatory Cascades in Streptomyces: Yield Enhancement and Metabolite Mining. Frontiers in Microbiology 11.

Yang, Z., Liu, C., Wang, Y., Chen, Y., Li, Q., Zhang, Y., Chen, Q., Ju, J., Ma, J., 2022. MGCEP 1.0: A Genetic-Engineered Marine-Derived Chassis Cell for a Scaled Heterologous Expression Platform of Microbial Bioactive Metabolites. ACS Synth. Biol. 11, 3772–3784. 10.1021/acssynbio.2c00362

Zaburannyi, N., Rabyk, M., Ostash, B., Fedorenko, V., Luzhetskyy, A., 2014. Insights into naturally minimised Streptomyces albus J1074 genome. BMC Genomics 15, 97. 10.1186/1471-2164-15-97

