## Supplementary material for "Characterization and Optimization of *Streptomyces albidoflavus* MD102 as a Heterologous Expression Chassis"

**Table S1.** Strains used in this study**.**

| Strain | Description |
|---|---|
| ***E. coli*** **TOP10** | For cloning |
| ***E. coli*** **ET12567/pUZ8002** | For conjugative transfer of plasmids into *Streptomyces spp.* |
| ***S. albidoflavus*** **MD102** | Isolated from Sungei Buloh Wetland Reserve, Singapore (1°26'41.172"N, 103°43'36.12"E); wild type genotype |
| ***S. albidoflavus*** **MD102Δ1** | ΔC16 |
| ***S. albidoflavus*** **MD102Δ2I1** | ΔC16 ΔC23a::ΦBT1-*attB* |
| ***S. albidoflavus*** **MD102Δ3I2** | ΔC16 ΔC23a::ΦBT1-*attB* ΔC5P19::*bldA* |
| ***S. albidoflavus*** **MD102Δ3I3** | ΔC16 ΔC23a::ΦBT1-*attB* ΔC5P19::*bldA* ΔC5P13::*gpps* |
| ***S. albidoflavus MD102SL01*** | ΔC16 ΔC23a::ΦBT1-*attB* ΔC5P19::*bldA* ΔC5P13::*gpps* ΔC23b ΔC23c ΔC1 |
| ***S. albidoflavus*** **MD102SL01::fur1** | *S. albidoflavus* MD102SL01 with pIJ8630-fur1 for heterologous production of flaviolin |
| ***S. coelicolor*** **M1154** | *Streptomyces* chassis strain |
| ***S. lividans*** **TK24** | *Streptomyces* chassis strain |

**Table S2.** Genes putatively involved in aromatic degradation found in *S. albidoflavus* MD102 but not in other common *Streptomyces* strains.

| Predicted function | Function | *S. albidoflavus* MD102 (PEG) | *S. albidoflavus* J1074 | *S. coelicolor* A3(2) | *S. avermitilis* MA-4680 |
|---|---|---|---|---|---|
| **Aromatic dioxygenation** | 2,3-dihydroxybiphenyl 1,2-dioxygenase (EC 1.13.11.39) | 589 | X | X | ✓ |
| **Biphenyl Degradation** | 2-hydroxy-6-oxo-6-phenylhexa-2,4-dienoate hydrolase (EC 3.7.1.-) | 590 | X | X | X |
| **Aromatic dioxygenase** | Cysteine dioxygenase (EC 1.13.11.20) | 1211 | ✓ | X | X |
| **Aromatic dioxygenation** | Cysteine dioxygenase (EC 1.13.11.20) | 2071 | ✓ | X | X |
| **Aromatic dioxygenation** | Quercetin 2,3-dioxygenase (EC 1.13.11.24) | 2946 | X | X | X |
| **Biphenyl Degradation** | 2-hydroxy-6-oxo-6-phenylhexa-2,4-dienoate hydrolase (EC 3.7.1.-) | 3330 | ✓ | X | X |
| **Biphenyl Degradation** | 2-keto-4-pentenoate hydratase (EC 4.2.1.80) | 6088 | X | X | X |
| **Biphenyl Degradation** | Acetaldehyde dehydrogenase, acetylating, (EC 1.2.1.10) in gene cluster for degradation of phenols, cresols, catechol | 6089 | X | X | X |
| **Biphenyl Degradation** | 4-hydroxy-2-oxovalerate aldolase (EC 4.1.3.39) | 6090 | X | X | X |
| **Biphenyl Degradation** | 2-hydroxy-6-oxo-6-phenylhexa-2,4-dienoate hydrolase (EC 3.7.1.-) | 6091 | X | X | X |
| **Aromatic dioxygenation** | 3-carboxyethylcatechol 2,3-dioxygenase (EC 1.13.11.16) | 6093 | X | X | X |

**Table S3.** Plasmids used in this study.

| Plasmid | Description | Reference |
|---|---|---|
| **pIJ12551** | Constitutive expression vector under the control of ermE*p | (Sherwood et al., 2013) |
| **pIJ8630** | EGFP reporter vector | (Sun et al., 1999) |
| **pIJ8630-ermEp*** | EGFP reporter vector under the control of ermEp* | This work |
| **pIJ8630-SF14p** | EGFP reporter vector under the control of SF14p | This work |
| **pIJ8630-kasOp*** | EGFP reporter vector under the control of kasOp* | This work |
| **pIJ8630-gapdhp(EL)** | EGFP reporter vector under the control of gapdhp(EL) | This work |
| **pIJ8630-gapdhp(KR)** | EGFP reporter vector under the control of gapdhp(KR) | This work |
| **pIJ8630-fur1** | Type III PKS *fur1* gene in pIJ8630 vector under the control of gapdhp(KR) | This work |
| **pIJ10702** | Cosmid vector derived from SuperCos1 | (Yanai et al., 2006) |
| **pSMART-BAC-S** | BAC-derived *E. coli-Streptomyces* shuttle vector | (Wu et al., 2012) |
| **pESAC13A** | PAC-derived *E. coli-Streptomyces* shuttle vector | (Sosio et al., 2000) |
| **pCRISPR-Cas9** | CRISPR-Cas9 vector for actinomycetes | (Tong et al., 2015) |
| **pCRISPR-Cas9-BGC16** | pCRISPR-Cas9 derivative for the knockout of BGC16 | This work |
| **pCRISPR-Cas9-BGC23a-ΦBT1-attB** | pCRISPR-Cas9 derivative for the knockout of BGC23a and insertion of *ΦBT1-attB* | This work |
| **pCRISPR-Cas9-BGC5P19-bldA** | pCRISPR-Cas9 derivative for the knockout of BGC5's P19 gene and insertion of *bldA* | This work |

| | | |
|---|---|---|
| **pCRISPR-Cas9-BGC5P13-gpps** | pCRISPR-Cas9 derivative for the knockout of BGC5's P13 gene and insertion of *gpps* | This work |
| **pCRISPR-Cas9-BGC23b** | pCRISPR-Cas9 derivative for the knockout of BGC23b | This work |

**Table S4.** The biosynthetic gene clusters in *S. albidoflavus* MD102 predicted by antiSMASH are shown, along with the subsequent genome modification efforts.

| Cluster No. | Locus | Most similar known cluster | Type [a] | In J1074 | In Post Modification Strain |
|---|---|---|---|---|---|
| 1 | 72006-147709 | Unknown Terpene-NRPS | Terpene, NRPS | No | Deleted |
| 2 | 301189-348721 | Alteramide | T1PKS, NRPS | Yes | Yes |
| 3 | 385976-411017 | Hopanoid | Terpene | Yes | Yes |
| 4 | 477446-485282 | Unknown Bacteriocin | RiPP-like | Yes | Yes |
| 5 | 733851-772692 | Paulomycin | PKS-like | Yes | Deleted::*bldA*, *gpp* |
| 6 | 943245-953524 | Unknown Bacteriocin | RiPP-like | Yes | Yes |
| 7 | 1196271-1256511 | Unknown NRPS | NRPS | Yes | Yes |
| 8 | 1352620-1366013 | Unknown Siderophore | NRPS | Yes | Yes |
| 9 | 1621886-1642836 | Geosmin | Terpene | Yes | Yes |
| 10 | 1955508-1975564 | Albaflavenone | Terpene | Yes | Yes |
| 11 | 2465198-2497664 | Thiopeptide | Thiopeptide | Yes | Yes |
| 12 | 2799470-2820002 | Lanthipeptide | Lanthipeptide | Yes | Yes |
| 13 | 3247182-3294215 | Butyrolactone | Type I PKS | No | Yes |
| 14 | 3586813-3636066 | Unknown NRPS | NRPS | Yes | Yes |
| 15 | 3889944-3994457 | Surugamide | NRPS | Yes | Yes |
| 16 | 4478518-4521430 | Unknown NRPS | NRPS | Yes | Deleted |
| 17 | 4746805-4758625 | Desferrioxamine B | NRPS | Yes | Yes |
| 18 | 5659455-5669853 | Ectoine | Ectoine | Yes | Yes |
| 19 | 6344976-6388218 | Indigoidine | NRPS | Yes | Yes |
| 20 | 6516019-6544745 | Isorenieratene | RiPP-like, Terpene | Yes | Yes |
| 21 | 6,633,062-6,644,429 | streptamidine | RiPP-like | Yes | Yes |
| 22 | 6659980-6701077 | Unknown PKS | Type III PKS | Yes | Yes |
| 23a | | Candicidin | Type I PKS | Yes | Deleted::ϕBT1 site |
| 23b | 6707464-6978266 | Antimycin | Type I PKS-NRPS | Yes | Deleted |
| 23c | | Unknown PKS | Type I PKS | Yes | Deleted |

[a] Biosynthetic Cluster Types: NRPS, non-ribosomal peptide synthetase; PKS, polyketide synthase

**Table S5.** Protein sequences of the biosynthetic enzymes encoded by the putative terpene-synthesizing BGC (i.e. BGC6) from *S. tasikensis* P46.

| Predicted function | Amino acid sequence |
|---|---|
| GGPPS | MTDAALDEFLDRKSLTAPSHQMLELVETLRGFLSSGGKRIRPVMCLCGWYAAGGKETPRP<br>VVKAAASLELFHACALIHDDVMDNSDARRGRLTLHRLLAERHRRRDPPGNAERFGTNAAI<br>LLGDLALAWSDEMFHTAGLSPAQVRAALPVLDAMRSEVMFGQYLDLLATGRPTGDVEE<br>ALMASRFKTAKYTVERPLHIGAALAGSGPAIRDALTAYALPVGEAFQLRDDLLGVFGDSR<br>QTGKPVDDDLREGKCTVLMALAVARADAAQLRVLRSLVGRSDLDAEEADAIRDVLVSTG<br>ARSVVDRMITSRCRRALSVLDRAPFPLPATNALRRLAHSASVRTS |
| Terpene synthetase | MFDIAPQRTGTSSPARHDEKAEGFFLPELPRLLPVAYHPKAAQIEFRSNAWLRRYLSGCFA<br>GEGELLKFLRERVGLYGPLIAPTADEQHALDLADFYHFVGVIDTMAADHSGLGASHCGAR<br>DVFDRIIADFAAGIEPGMSDNSPFGPAARDLWLRISAGLTPHQVERFRTSMSSFLRGVASEL<br>PYQLNGSVPDYDTYMAVRRDSFGCDFILLLTEYSLAVDMTELAASPQFAKVHAHAMRQLI<br>LVNDVLSLRKELGDPMNAVRVLRRHNGLTLQQAVDAVCELAERHERAYIAARDAVRYG<br>PFGAHTDVRTYLEGLDHLLAGSQEYEYLTPRYFGDGSVWDGSTSGWISLTAPIARFLPEAG<br>PTSEQRRTKAVRIHTRRS |
| Cytochrome P450 | MTTAHEARAESCPSTRGTAPGGLPLLGHALPLRRRPLEFLAALPAQGDLVEVRLGPRRMY<br>LACHPDLVQQVLRDSRTFDKGGPMFDKVRLLTGNGLATSCWAEHRRQRRLVQPAFHQER<br>MAGYAAVMDHEITSMLDSWQEGRVLDVHAAMQALTARVIVRTLFSTRIEDWAVDEIQRC<br>LPVITRGFFTRMVAPLGLVQKLPTRSNREFDRALERMRNVIDQTVSSHPTGTDHGDLLSGL<br>LRAEDEETGERLAGHEIHDQVMTLLMGAIETTSNTLAWTYHLLGENPEAEARLHREIDSVL<br>PARRPGFDDLPHLGYTQRVVTEALRLYPPTWLLTRSTTCEAELAGLRLAPGTTVALSFYAL<br>GHNPALFCDPERFDPDRWLPERAKTVPRGAWNSFGGGSRKCIGDRFATIETTLVLAAVAS<br>GWRLKPHPGPSIRPEPKASLSTGPLPMIPERRKTSGAGAPAARSPWTLTAPPPDDHC |

REFERENCES


Sherwood, E.J., Hesketh, A.R., Bibb, M.J., 2013. Cloning and Analysis of the Planosporicin Lantibiotic Biosynthetic Gene Cluster of Planomonospora alba. J Bacteriol 195, 2309–2321. https://doi.org/10.1128/JB.02291-12

Sosio, M., Giusino, F., Cappellano, C., Bossi, E., Puglia, A.M., Donadio, S., 2000. Artificial chromosomes for antibiotic-producing actinomycetes. Nat Biotechnol 18, 343–345. https://doi.org/10.1038/73810

Sun, J., Kelemen, G.H., Fernández-Abalos, J.M., Bibb, M.J., 1999. Green fluorescent protein as a reporter for spatial and temporal gene expression in Streptomyces coelicolor A3(2)This paper is dedicated to the memory of Kathy Kendrick, whose devotion to understanding the biology of Streptomyces was unsurpassed. Microbiology 145, 2221–2227. https://doi.org/10.1099/00221287-145-9-2221

Tong, Y., Charusanti, P., Zhang, L., Weber, T., Lee, S.Y., 2015. CRISPR-Cas9 Based Engineering of Actinomycetal Genomes. ACS Synth Biol 4, 1020–1029. https://doi.org/10.1021/acssynbio.5b00038

Wu, C.-C., Liles, M., Kakirde, K., Ye, R., Wagner, M., Krerowicz, A., Staley, M., Jasinovica, S., Drinkwater, C., Godiska, R., Mead, D., 2012. Next-generation functional and structural soil metagenomics.

Yanai, K., Murakami, T., Bibb, M., 2006. Amplification of the entire kanamycin biosynthetic gene cluster during empirical strain improvement of Streptomyces kanamyceticus. Proc Natl Acad Sci U S A 103, 9661–9666. https://doi.org/10.1073/pnas.0603251103
